# Sequence-free landscape inference for directed evolution

**DOI:** 10.1101/2025.10.08.681123

**Authors:** Sebastian Towers, Jessica James, Harrison Steel, Idris Kempf

**Affiliations:** Department of Engineering Science, University of Oxford, Parks Road, Oxford OX1 3PJ, UK

## Abstract

Directed evolution is a method for engineering biological systems or components, such as proteins, wherein desired traits are optimised through iterative rounds of mutagenesis and selection of fit variants. The process of protein directed evolution can be envisaged as navigation over high-dimensional landscapes with numerous local maxima. The performance of any strategy in navigating such a landscape is dependent on the ruggedness of that landscape. However, this information is generally unavailable at the outset of an experiment, and cannot currently be computed using analytical methods. Here we propose **SLIDE, S**equence-free **L**andscape **I**nference for **D**irected **E**volution, which consists of two parts. First, SLIDE provides an estimation for landscape ruggedness from a mutating population using only population-level phenotypic data and an estimation of mutation rate. Ruggedness information in itself is valuable in protein design, for instance in predicting evolutionary stability. Second, SLIDE offers a framework for using the estimated ruggedness metric to select high-performing parameters for directed evolution control. Using theoretical *NK* landscapes and four real-world protein fitness landscapes, we demonstrate improvement upon the performance of standard selection strategies, particularly on rugged landscapes, using a pipeline that could also be combined with emerging AI-based methods for driving direction evolution.

## Introduction

Fitness landscapes map DNA or amino acid sequences to a measure of fitness such as enzyme activity, cell growth, binding affinity, or stability [1]. The shape of a fitness landscape, be it smooth or rugged, has significant consequences for evolution and stability [2]. Inferring the ruggedness of a fitness landscape unlocks the ability to assess the robustness of a protein to mutations and to optimise directed evolution (DE), which is the focus of this work.

Various metrics have been proposed to define and quantify the ruggedness of fitness landscapes [3– 7]. Although these metrics have been shown to correlate across several empirical landscapes [7], most of them require full genotypic information to compute a ruggedness measure. An exception to this comes from spectral landscape theory [8], which describes fitness landscapes using tools of Fourier analysis. In the same way that a signal can be decomposed into a sum of waves, a fitness landscape can be decomposed into a sum of landscapes of different frequencies (ruggedness). Where low frequency terms dominate, the landscape can appear smooth, whereas where high frequency terms dominate, the landscape can appear rugged.

In this paper, we develop Sequence-Free Landscape Inference for Directed Evolution (SLIDE) to efficiently estimate the ruggedness of a landscape using phenotypic information only (Fig 1). Based on spectral landscape theory, we define a novel ruggedness metric related to the dominant Fourier terms, and show that this metric can be estimated from the average fitness decay of a population that is randomly mutated from a starting point. On smooth landscapes, the average fitness decays more slowly than on rugged landscapes. This phenomenon can be attributed to fitness correlations between variants, as random mutations are less likely to result in large fitness differences. Most importantly, the fitness decay curve can be acquired without large-scale sequencing data; instead it uses population-level fitness readouts and a mutation rate estimation.

**Figure 1.**
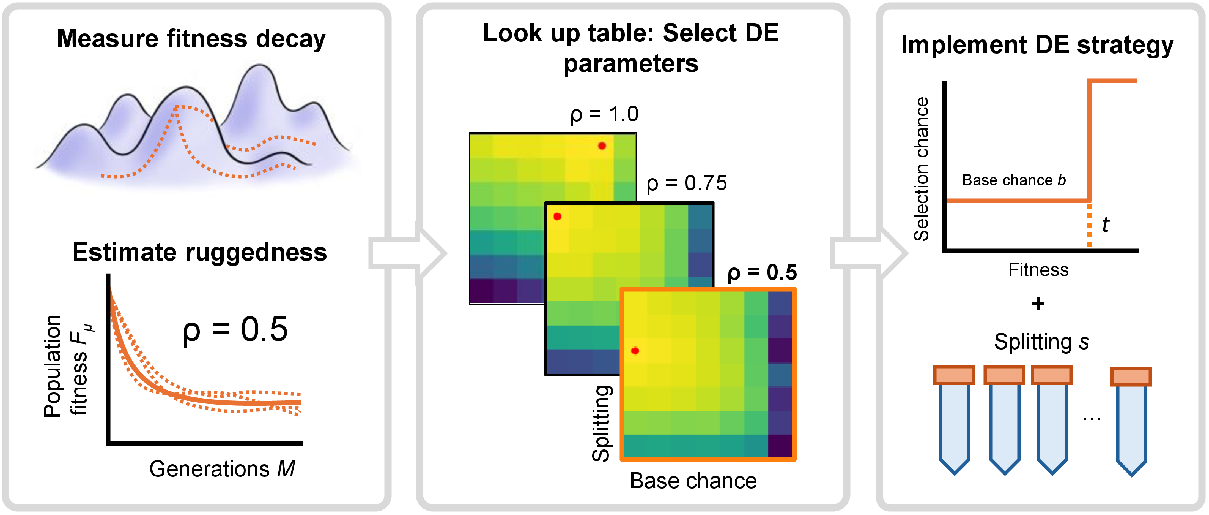
Schematic outline of a future DE pipeline. The initial phase of SLIDE involves random mutation away from a starting point, and measurement of the resulting fitness decay curve, which can be used to estimate the decay rate *ρ* as a measure of ruggedness. A look-up table matches *ρ* values of NK landscapes to optimal DE strategy parameters. This look-up table can be used to select appropriate parameters for the measured *ρ*. Parameters explored in this paper include base chance (applying a constant chance of selection that is irrespective of fitness) and population splitting [9].

In SLIDE, we combine ruggedness inference with DE. In DE, biological components such as proteins are engineered and improved through iterative rounds of selection and mutagenesis [10]. It can be seen as navigating a high-dimensional fitness landscape, aiming to guide the fitness of variants to a high-performing location on the landscape. Early DE optimisation techniques such as ProSAR used statistical models to correlate sequence with measured activity and iteratively design variants that maximize predicted fitness [11]. Since then, advances in machine learning have led to the development of more sophisticated algorithms that actively design the sequence of each variant that comprises the next generation [12–14]. Although effective, these methods are limited by their need for high-throughput sequencing data. Without sequencing data, most DE experiments simply select the top proportion of variants in each iteration [15–17], which will henceforth be referred to as the “baseline strategy”. This approach results in minimal landscape exploration, and is hence prone to getting trapped in local optima [9].

To address this limitation, several methods have been proposed for balancing landscape exploration and exploitation. Such methods include applying non-constant selection pressure [18], increasing the proportion of population selected [19], splitting the population into sub-populations, and including a base chance of selection that is irrespective of fitness [9]. However, effective application of these methods depends on knowledge of landscape properties such as ruggedness [9]. Inferring the ruggedness beforehand is essential for selecting strategy parameters to ensure the balance of exploration and exploitation. Here, we select these parameters based on a ruggedness estimation through SLIDE, where ruggedness and strategy parameters are identified prior to DE experiments. By doing so, SLIDE increases the likelihood of reaching higher-fitness variants within a fixed number of generations.

Any method for DE – both with and without sequencing – can integrate SLIDE to tune DE strategy parameters. In this paper, SLIDE is interfaced with strategies developed in our previous work [9], controlling base chance and population-splitting parameters. We evaluate our method for DE *in silico* on theoretical landscapes obtained from the *NK* model [20] and on four empirical landscapes [2, 21–23]. We show that SLIDE consistently finds the optimal parameters on *NK* landscapes, and outperforms or matches the baseline strategy on all empirical landscapes.

## Results

### Ruggedness inference from fitness decay rate

Ruggedness of a fitness landscape can be defined in terms of various mathematically distinct metrics (see [3–7]), which aim to capture how fitness changes when the landscape is traversed. Fig 2A-B shows examples of smooth and rugged landscapes for a protein of length *N* = 2 with 16 genotypes, where *σ*_*i*_ is used to denote the allele at gene locus *i* = {1, 2} (i.e. the set of possible amino acids, here *A* = 4). Intuitively, smooth landscapes should have high fitness variants clustered together, leading to high fitness correlations and relatively few local maxima. In contrast, rugged landscapes should exhibit large fitness differences between adjacent nodes, resulting in more local maxima and lower fitness correlations.

**Figure 2.**
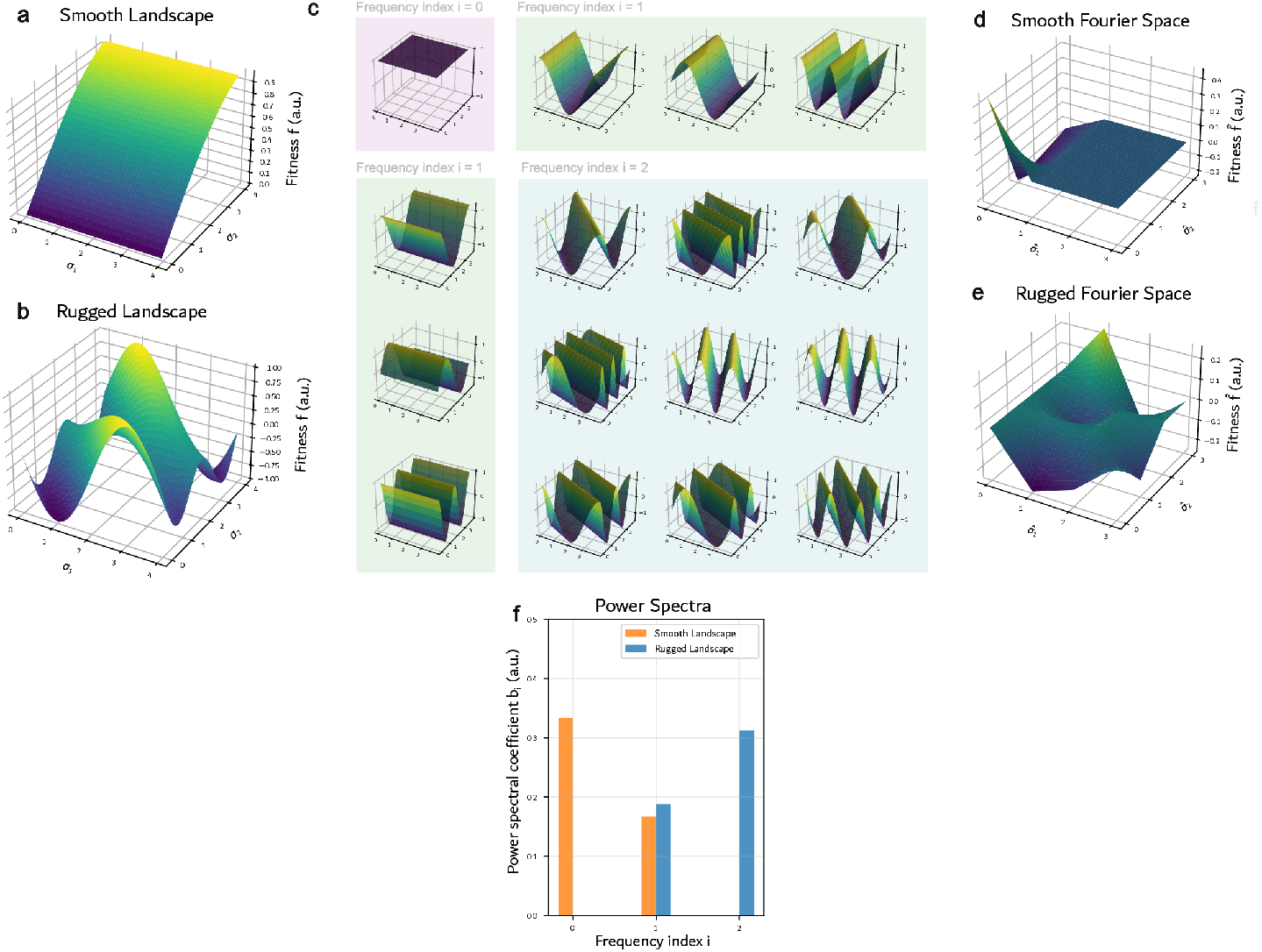
Demonstration of how two example landscapes can be transformed into a power spectrum. Examples of **A**: a smooth and **B**: a rugged landscape. **C**: Example Fourier basis. **D-E**: Smooth and rugged landscapes transformed into Fourier space with the example basis. **F**: The power spectrum, showing how the rugged landscape has a higher contribution from rugged terms compared to the smooth landscape.

In this work, we develop a metric to measure this behaviour based on spectral landscape theory [8], which decomposes landscapes into low-frequency (smooth) and high-frequency (rugged) components using a Graph Fourier transform (GFT). Analogous to a standard Fourier transform, the GFT decomposes the original landscape ***f*** to express it as a mixture of basis landscapes of increasing frequency, such as those shown in Fig 2C. For a protein of length *N* and *A* choices of amino acids, there are *A*^*N*^ basis landscapes grouped into *N* + 1 frequencies *λ*_0_, …, *λ*_*N*_. The weights of each basis landscape in ***f*** can be arranged to form a new landscape 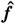 in Fourier domain, such as shown in Fig 2D-E. In the Fourier domain, the alleles *σ*_*i*_ are replaced with the Fourier components 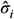 and the genotype fitness with the weight of the Fourier component 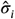. The information from Fig 2D-E can be further condensed by grouping the Fourier component weights for each frequency *λ*_*i*_ to yield the power spectrum, i.e. by computing the power *b*_*i*_ of the landscape over frequency *λ*_*i*_ (Fig 2F).

Based on the power spectrum from Fig 2F, we propose to extract a single, scalar ruggedness metric *ρ* as the power-weighted frequency average

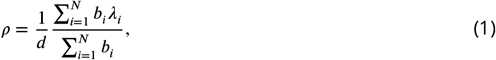

where *d* = *N*(*A* − 1) is a factor determined by landscape size/dimension, *b*_*i*_ the power spectral coefficient at frequency index *i*, and *λ*_*i*_ = *Ai* a frequency-specific parameter (see Methods). The quantity in equation (1) represents the center of mass of the spectrum, and can be related to the *R*^2^ ruggedness measure from [7]. It is unique because it is a robust ruggedness metric that can be easily computed from phenotypical data extracted from a randomly mutagenised population. In Methods, we show that if each population member has accumulated an average of *μ* single-point mutations, the fitness decay and the ruggedness metric from equation (1) are related by:

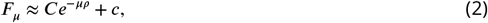

where *F*_*μ*_ is the population’s average fitness, *c* is the average landscape fitness, and *C* is a constant that depends on the initial population and can be negative if the intial population fitness is lower than *c*. A similar relationship, independent of the initial population, can be obtained by squaring *F*_*μ*_ and averaging over different initial populations (see Methods). In both cases, the decay rate *ρ* can be interpreted as the ruggedness of the landscape by relating it to the different Fourier components from Fig 2: for a large *ρ*, higher-frequency terms dominate the landscape, and for a small *ρ*, lower-frequency terms dominate. In certain analytically tractable models, this link becomes explicit. For example, in *NK* landscapes, the epistatic interaction parameter *K* and *ρ* are related by *ρ* ≈ (*K* + 1)*/N* (see equation (S2) in the Supplementary). On more rugged landscapes, the decay rate *ρ* will be larger than on smoother landscapes, and the mean population fitness will converge more quickly to the landscape average. This observation is the essential phenomena that our work exploits to estimate the landscape ruggedness through measurements of the mean fitness of a population as it accumulates mutations. In other words, suppose that each population member is mutagenised with a single-point mutation on average, and the mean fitness recorded as *F*_1_, *F*_2_, …, *F*_*M*_ over *M* generations. These values can be used to estimate the parameters *C, ρ*, and *c* in equation (2). This is the first part of our two-step procedure referred to as SLIDE.

The trend from equation (2) is illustrated in Fig 3a, which shows the mean population fitness decay from mutating populations on two *NK* landscapes of differing ruggedness, and the approximation from equation (2). Fig 3a shows that on landscapes with greater ruggedness ((*K* + 1)*/N* = 0.75), the rate of fitness decay is higher than that of the smooth landscape ((*K* + 1)*/N* = 0.1). This is further validated in Fig 3b, which shows that the estimated *ρ* corrlates strongly with (*K* + 1)*/N* ∈ {0.1, 0.2, …, 1}.

**Figure 3.**
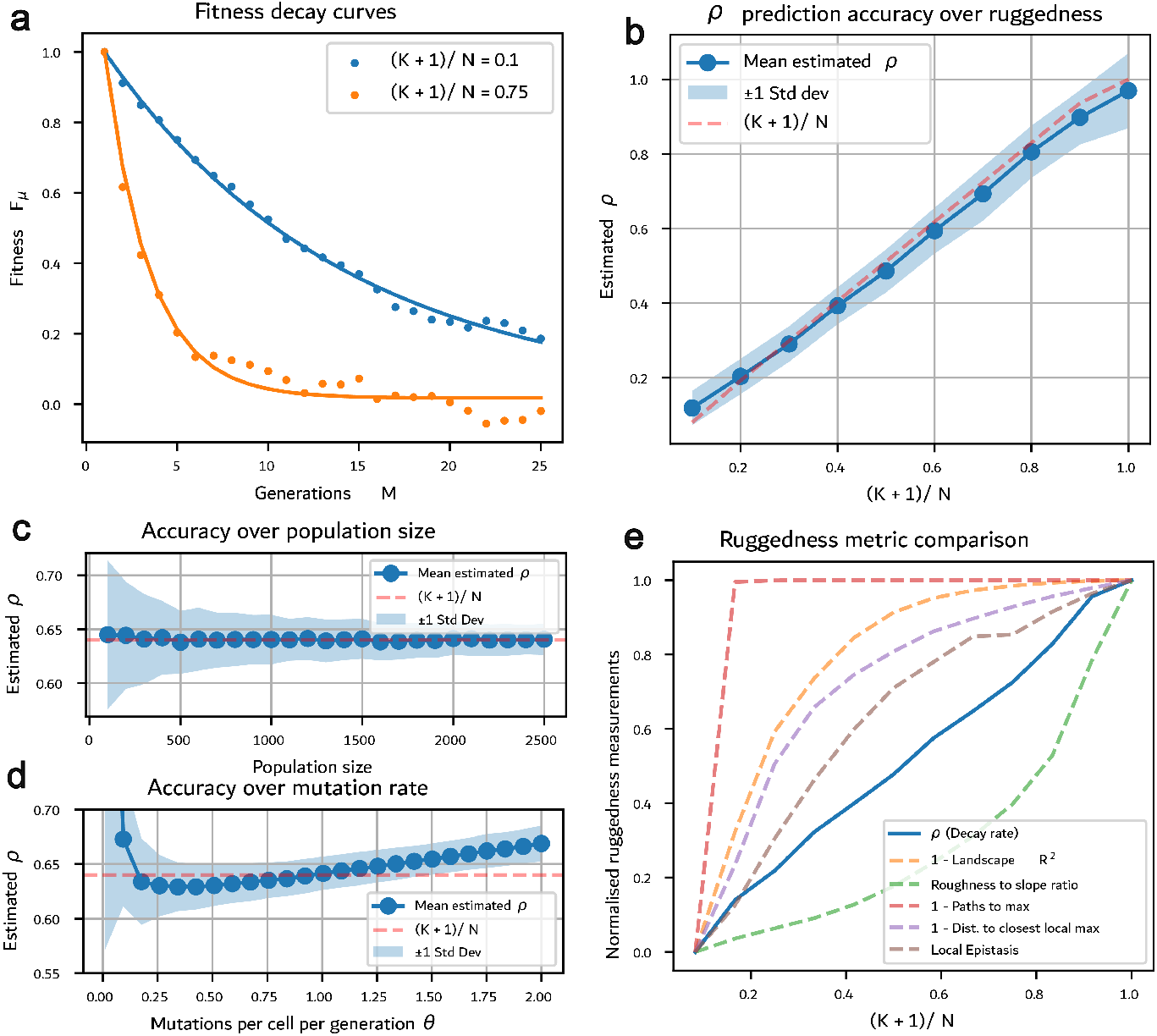
Fitness decay rate of a randomly mutating population, as a metric for landscape ruggedness. **A:** Simulations of two populations on two *NK* landscapes ((*K* + 1)*/N* = 0.1 and 0.75, respectively, with N = 20) mutating away from a starting point (population size = 300, mutation rate *θ* = 0.5 mutations per generation). Exponential fits to equation 1 are indicated by solid lines. **B:** Estimated *ρ* values across landscapes of increasing ruggedness using 25 generations and a mutation rate of 0.5 per generation. The figure shows mean and standard deviation for 300 estimates. **C–D:** Accuracy of *ρ* over increasing population size (C) and mutation rate (D) for 500 estimates. **E:** Comparison of ruggedness metrics on 600 *NK* landscapes of increasing ruggedness ((*K* + 1)*/N*). All metrics are normalised between 0 (least rugged) and 1 (most rugged).

Due to the noise associated with observing a randomly mutating population, the reliability of *ρ* estimates also depends on the parameters of the observed population, including the population size and the mutation rate. Fig 3c shows *ρ* estimates over increasing population sizes. For each population size, *ρ* is estimated 500 times with Fig 3 showing mean and standard deviation (Std Dev). For small population sizes, the ruggedness estimate exhibits higher variability due to noise in the mean population fitness. For population sizes exceeding 2,000, the standard deviation is less than 5 % of the “true” decay rate. The effect of mutation rate is analysed in Fig 3d, which shows mean and standard deviation of *ρ* for an increasing mutation rate *θ*. For small *θ*, only a small fraction of the fitness decay curve in Fig 3a is captured, leading to a large variability of the estimate from equation (2). For very large *θ*, the mean population fitness decays more rapidly to the landscape average, again leading to a large variability of the estimate from equation (2).

To further validate equation (2) and its characteristic decay rate *ρ*, Fig 3e provides a comparison between *ρ* and other ruggedness measures across *NK* landscapes (See Supplementary section 3). The other ruggedness measures are the landscape *R*^2^ measure [7], the roughness to slope ratio [3], number of paths to the global maximum from the opposite position on the landscape [6], the distance to the nearest local maximum [5], and the local epistasis metric [7]. Figure 3e shows that although all metrics increase with (*K* + 1)*/N*, they express either concave (roughness to slope ratio), linear (decay rate), or convex curves (all other metrics). Because all metrics increase with *K*, this allows one to compare the ordering (w.r.t. ruggedness) of landscapes across metrics, but the differences in curve shapes prohibits comparison of distances (i.e. quantitative ruggedness differences) between landscapes across metrics. However, one can mathematically prove the decay rate *ρ* directly correlates with (*K* + 1)*/N* (see equation (S2) in the Supplementary Material), and the same relationship holds regardless of *N*. Overall we believe this provides a second benefit of our methodology - that it provides an interpretable measure of ruggedness that has immediate meaning in the widely-studied *NK* landscapes.

### Ruggedness inference on real-world landscapes

The performance of SLIDE on real-world empirical landscapes was assessed by testing it on four experimentally-measured landscapes: GB1 (4 sites) [2], ParD3 (3 sites) [21], TrpB (4 sites) [22] and TEV (4 sites) [23]. It was found that the variability over *ρ* estimations via *F*_*μ*_ from equation (2) was significantly higher than on *NK* landscapes, on account of *NK* landscapes being much more homogenous over the space than the empirical ones, i.e. landscape properties computed from a small set of local measurements are more representative of global landscape properties. To account for this effect, it is therefore necessary to estimate *ρ* via 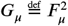 from equation (2), which (as per Methods) requires to sample, square, and average fitness decays from multiple starting points. Despite the added labour required to measure multiple starting points, such a set up remains scalable as it requires mutagenesis only, something which is commonly conducted in a pooled format for library generation. For each landscape, this method is used to compute a “true”, global decay rate, obtained from averaging fitness decays from 10,000 starting points (dashed lines in Fig 4a). The true decay rates are compared against estimates that use an increasing number of averages. Interestingly, the variability over *ρ* differs significantly between landscapes, with TEV in particular exhibiting over 4× more variability than GB1 over 1,000 samples.

**Figure 4.**
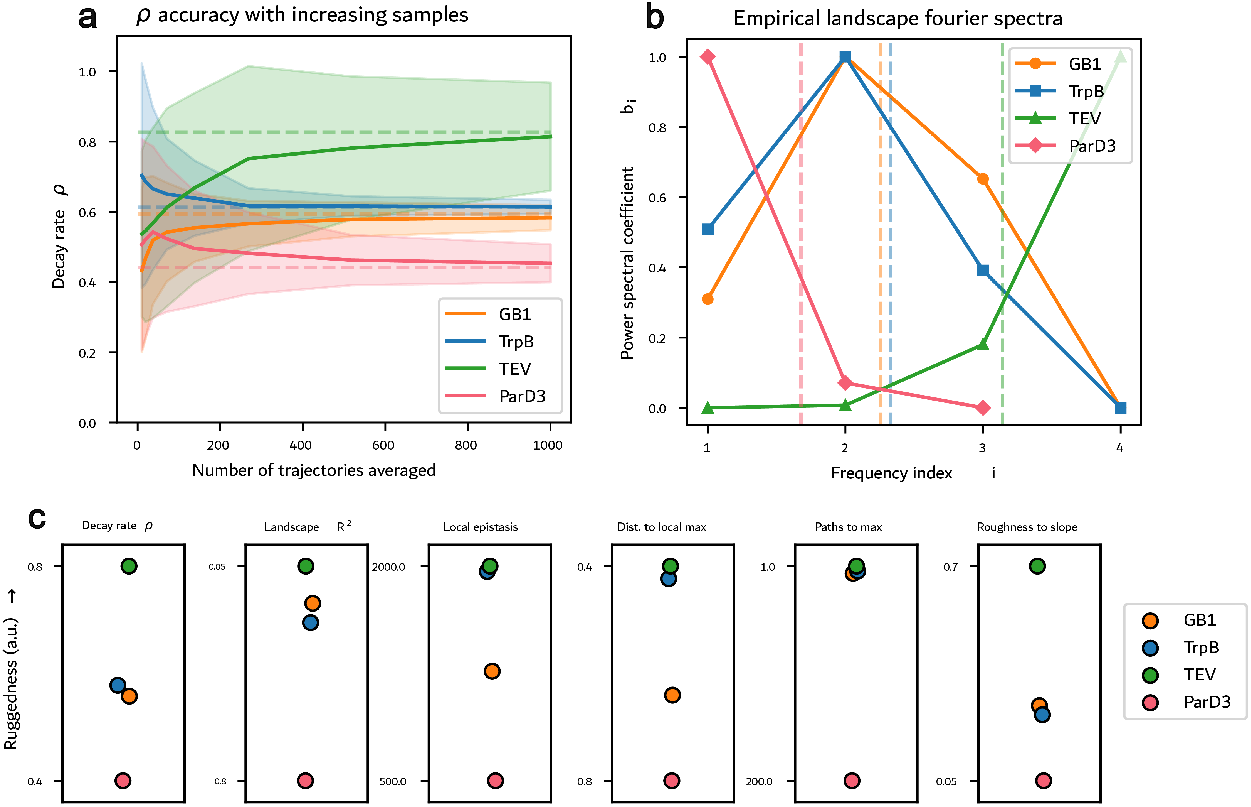
Ruggedness estimation on empirical landscapes. **A:** Estimations of *ρ* with increasing numbers of starting points. Starting points were sampled uniformly across the space, with mutation rate 0.1 per generation, population size = 2500. **B:** Fourier spectra of each complete landscape computed using GFT (Methods). Estimated *ρ* value indicated by dashed line. **C:** Comparison of ruggedness metrics applied to the empirical landscapes.

The *ρ* values, as estimated by the decay rates, were then compared to the computed Fourier spectra of the complete landscapes (Fig 4b). It can be seen that the resulting *ρ* values are broadly equivalent to the center of mass of the Fourier terms, providing empirical justification for the relationship between fitness decay rates and Fourier spectra.

Fig 4c compares the different ruggedness measures across the empirical landscapes. Given the variation in scaling and linearity between metrics, comparisons should be focused on the ordering of landscapes within each metric, as opposed to comparisons of raw values between metrics. Local epistasis, distance to local maximum, and paths to global maximum agree with *ρ* in terms of their ordering of all the empirical landscapes tested. Landscape *R*^2^ and roughness to slope ratio agree with the extremities (TEV as the most rugged, ParD3 as the least), however place GB1 and TrpB in a different order. Similarly, our metric *ρ* groups together TrpB and GB1 landscapes as having a similar, intermediate ruggedness, whereas other methods separate these landscapes, or locate them very close to the TEV landscape. Given that other ruggedness metrics generally agree with the ordering of *ρ* on empirical landscapes provides increased confidence in the metric, which has the major advantage over all others of being experimentally measurable sequencing data (see Supplementary section 6 for a biological interpretation of these ruggedness values).

### Optimising DE outcomes with SLIDE

Inferring the ruggedness of a fitness landscape prior to DE allows informed decisions to be made about the DE strategy. In our previous work [9], it was shown that increased population splitting and base chance – a constant probability of selection that is irrespective of fitness – can improve performance on rugged landscapes, but reduce performance on smooth landscapes. By utilising the relationship between ruggedness and fitness decay rate demonstrated in this work, it is therefore possible to optimise DE outcomes without the need for sequencing.

In order to explore the relationship between decay rate and optimal strategy, a large parameter sweep was conducted over 100 different *NK* landscapes at fixed mutation rate and population size (Supplementary section 2). For each landscape, 14,700 DE simulations were conducted over 49 strategy combinations. Fig 5a (dotted line) displays the optimal base chance and splitting values over landscapes of increasing epistatic parameter *K*. As *ρ* ≈ (*K* + 1)*/N*, this data can be used as a SLIDE look-up table to select an appropriate combination of DE parameters from *ρ*. To demonstrate SLIDE on NK, for each of the 100 landscapes, 250 decay curves were simulated. From these curves, decay rate was measured and used to predict optimal parameters, the outcome of which is also shown in Fig 5a (error bars).

**Figure 5.**
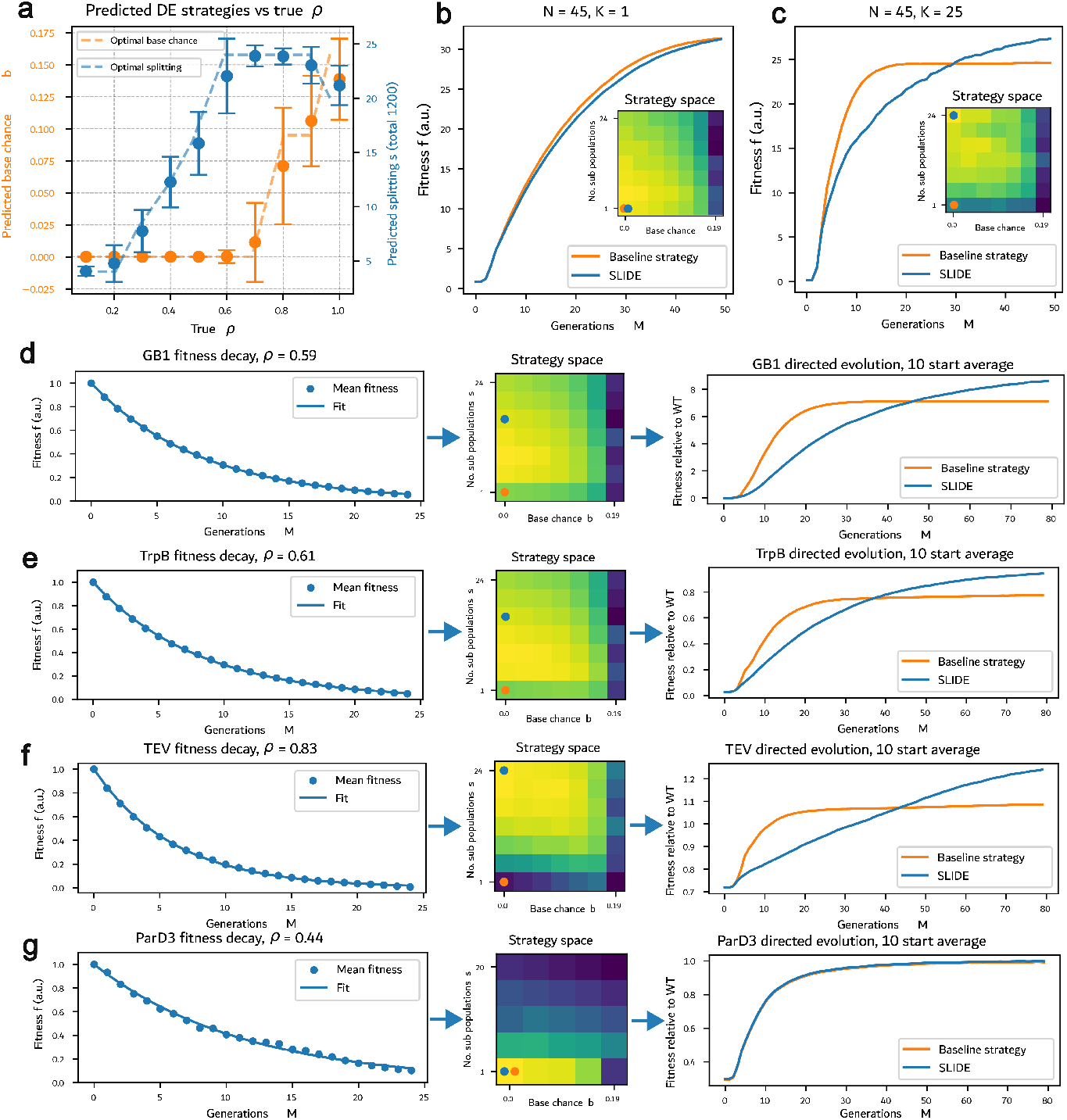
DE using SLIDE on *NK* and empirical landscapes. **A:** Representation of the SLIDE look up table for *NK*. Generated from 100 *NK* landscapes of increasing ruggedness, with 14,700 simulations in total over 49 strategy combinations. Base chance options = {0.00, 0.03, 0.06, 0.10, 0.13, 0.16, 0.19}, splitting options = {24, 20, 16, 12, 8, 4, 1}. Error bars represent the accuracy of SLIDE in selecting the true optimal parameters, from 250 simulations. **B-C:** Implementations of SLIDE on an example smooth and rugged *NK* landscape, respectively. Population size = 1200, mutations = 0.1/N per generation. Implementations of SLIDE on **D:** GB1, **E:** TrpB, and **F:** TEV. Decay curves from 10,000 starts, mutations = 0.025 per generation, population size = 2500. SLIDE strategy look-up table generated for A = 20, N = 4. DE simulations conducted from 10 random starts, population size = 1200, mutations = 0.01 per generation. **G:** ParD3. Parameters as above, except base chance options = [0.00, 0.05, 0.10, 0.14, 0.19], splitting options = [20,15,10,5,1]. SLIDE strategy look-up table generated for *A* = 20, *N* = 3. DE population size = 60.

Two examples of DE curves on *NK* landscapes are displayed in Fig 5b–c. Fig 5b displays a smooth landscape (*ρ* ≈ 0.02), where the strategy selected by SLIDE is highly similar to the baseline strategy (which is already near the optimum). Fig 5c, however, displays a more rugged landscape (*ρ* ≈ 0.55), on which the baseline strategy is highly sub-optimal and gets trapped at a local optimum within 10 generations. SLIDE, on the other hand, selects the optimal strategy within the strategy space and achieves > 15 % higher fitness than the baseline strategy after 50 generations.

A similar approach was then applied to the four combinatorially-complete empirical landscapes GB1, TrpB, TEV and ParD3 (Fig 5d-g). The strategy look up, as displayed in Fig 5a, was re-generated using *NK* landscapes of the same dimensions as the landscape: *A* = 20, *N* = 4 (or *N* = 3 for ParD3). As well as this, it was necessary to increase the number of starting points sampled to generate an accurate estimation (as described in the previous section). SLIDE out-performed the baseline strategy on all examples, apart from ParD3, for which the baseline strategy was optimal (and therefore SLIDE selected the same parameters). On GB1, TrpB and TEV, it achieved relative fitness gains of 17.4 %, 18.2 %, and 29.5 % fitness improvement respectively over the baseline approach, which commonly gets trapped at local maxima.

## Discussion

In this paper, we showed that the average fitness decay of a population accumulating mutations in the absence of selection pressure can be linked to the power spectrum of the underlying fitness landscape. Specifically, we demonstrated that the average fitness decay can be approximated by an exponential function, with the decay rate related to the dominant frequency components of the landscape’s Fourier spectrum. This decay rate serves as a practical ruggedness metric, which we compared with established metrics on *NK* models and empirical protein landscapes.

Building on our ruggedness metric, we introduced SLIDE (Sequence-Free Landscape Inference for Directed Evolution), a framework that estimates landscape ruggedness from average fitness decay curves. Unlike existing methods, SLIDE requires only phenotypic data—removing the need for large-scale sequence-level information. As long as fitness can be coupled to a measurable phenotype such as fluorescence, SLIDE enables ruggedness estimation from bulk population-level measurements and knowledge of the average mutation rate. This feature makes SLIDE uniquely scalable and accessible for experimental use. We validated SLIDE on both synthetic *NK* landscapes and empirical datasets, showing that it can result in up to a 29.5 % improvement in directed evo-lution performance compared to the most common baseline directed evolution strategy. While this paper applied directed evolution strategies from [9], the ruggedness metric introduced by SLIDE is broadly compatible with a wide range of advanced directed evolution approaches, including AI-driven workflows. For example, in active learning-assisted directed evolution [13], machine learning techniques are used to guide the selection, balancing exploration of unknown phenotypes with exploitation of high-performing variants. Here, the ruggedness metric obtained from SLIDE could be used to modulate the balance of exploration and exploitation. Similarly, in cluster learning-assisted directed evolution [24], SLIDE could be used to tailor clustering granularity.

Although this work focused primarily on the theoretical and computational aspects of SLIDE, the framework is designed to be experimentally implementable. The key requirement is a biological protocol for introducing mutations without applying selection pressure. One example is random *in vitro* mutagenesis, which allows targeted exploration of specific regions of the genotype space. Mutant libraries generated in this way can be expressed in cells, with fitness linked to a measurable phenotype like fluorescence. By iteratively measuring average fluorescence across generations and estimating mutation accumulation (or by simultaneously producing samples with increasing levels of mutation), it becomes possible to construct fitness decay curves and apply SLIDE in a laboratory setting. Our simulations suggest that even modest experimental setups—using populations of 100 variants over fewer than 10 mutagenesis iterations—could yield reliable decay estimates. This opens the door to practical, low-cost ruggedness estimation across diverse protein families and fitness landscapes.

The requirement of a selection-free environment could be removed by extending the mathematical framework developed in Section Methods. The selection pressure (or any other selection rules) could be factored into the mutagenesis model and the ruggedness estimation algorithm adapted. This would offer new possibilities for experimental ruggedness estimation, such as continuous *in vivo* mutagenesis [25] in small-scale bioreactors. Additionally, it would allow SLIDE *during* directed evolution experiments as an online feedback signal.

In summary, SLIDE offers a novel, sequence-free method for estimating the ruggedness of protein fitness landscapes using average phenotypic data. These ruggedness estimates can inform experimental strategies in directed evolution, helping to tailor protocols to the underlying structure of the landscape. Next to directed evolution, SLIDE’s ruggedness metric could be extended to characterise more complex landscapes, such as those underlying synthetic biological circuits. Large-scale mapping of ruggendess in such systems could provide a quantitative basis for designing circuits with greater genetic stability and functional robustness, complementing recent efforts to understand and engineer the evolutionary potential of synthetic constructs [26].

## Methods

### Decay Rate Metric

To show why the fitness decay rate *ρ* in Eq (2) can be used as a ruggedness metric, we give a brief overview of spectral landscape theory. We will represent a population accumulating single-point mutations as a stochastic signal in a graph, modelling mutagenesis and genotype space. For a gene of length *N*, we denote the genotype space by 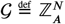 (the space of positive integers up to *A*, in a vector of *N* entries), containing |𝒢|= *A*^*N*^ genotypes 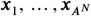. Each genotype can be written As 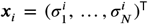, where 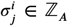 is the allele at locus *j*. For example, if *A* = 20 represents the number of possible amino acids at each position in a protein of length *N*, the landscape consists of *A*^*N*^ distinct genotypes. These genotypes can be represented as nodes in a graph, with each node being assigned a fitness value 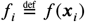, where *f* : 𝒢 ↦ ℝ characterises the fitness landscape. A population is represented by 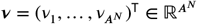, where each ν gives the frequency of genotype ***x***_*i*_. Equivalently, ***ν*** can be defined as a probability distribution, in which case ∑_*i*_ ν_*i*_ = 1 with all ν_*i*_ ∈ [0, 1]. The mean population fitness is then computed as *F* = ***f*** ^T^ν = ∑_*i*_ *f*_*i*_ν_*i*_, where 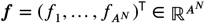 is the stacked vector of fitness values.

The edges of the graph connect genotypes that differ by a single mutation, so that every node is connected to 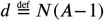 other nodes, which is also referred to as a Hamming graph [27]. For node transitions, we define the single-mutation operator 𝒟 to map an initial population ***ν*** to 𝒟 (***ν***), where every population member has undergone a single, random, mutation (see the Supplementary section 1 for details on our mutation model). It can be shown that 𝒟(***ν***) = *D****ν***, where 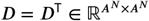 is the adjacency matrix of our graph multiplied by 1*/d*. We assume that the number of mutations *X*_*M*_ that occurs between two observations is Poisson distributed with mean *μ*, so that the population is expected to evolve as

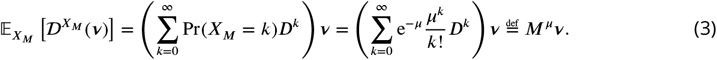

Our mutation operator *M*^*μ*^ can be understood as the transition function of a continuous time Markov chain with a single genotype randomly mutating at rate *μ*. It can be simplified using matrix exponentials as

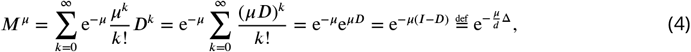

where we substituted the graph Laplacian 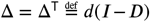. The mean fitness *F*_*μ*_ of our population after an average of *μ* mutations is therefore

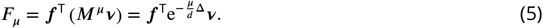

If necessary, the dependency on the initial population ***ν*** can be omitted by taking the expectation of Eq (5) over many random starting populations. In particular, if ***ν*** is sampled uniformly from single-genotype populations, so that 𝔼[***νν***^T^] = *I/A*^*N*^, we can compute 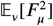 as:

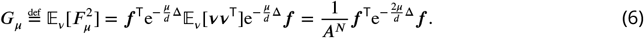

Eqs (5) and (6) can be further simplified by considering the special structure of a Hamming graph. Its real-valued, symmetric Laplacian admits a Fourier basis as eigenbasis [28], so that Δ can be factorised as Δ = *P* Λ*P* ^∗^, where 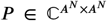 satisfying *P* ^∗^*P* = *I* contains the eigenvectors, and 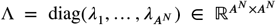, with the eigenvalues ordered as 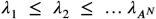. Analogous to the standard Fourier transform, the eigenvalues capture a frequency and the columns of *P* eigenfunctions, each of which can be interpreted as eigenlandscapes of increasing ruggedness for increasing *λ*_*i*_. For the Hamming graph Laplacian, there are *N* +1 unique eigenvalues with value *λ*_*i*_ = *Ai* and multiplicity 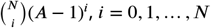 [29]. The matrix *P* may be chosen as 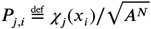, where 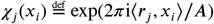 is the Fourier basis function, *r*_*j*_ ∈ 𝒢 the frequency index, and ⟨*r*_*j*_, *x*_*i*_⟩ the dot product defined on 𝒢 (dot product modulo *A*) [28]. For *λ*_0_ = 0, the eigenlandscape is a constant and for *λ*_1_ = *A*, the landscape is linear, i.e. depends only on the allele *σ*_*j*_ at a single locus *j* (Fig 2c). For larger eigenvalues, the eigenlandscapes are rugged, i.e. they depend on the allele at multiple loci (exactly *i* different loci for *λ*_*i*_ = *Ai*).

The special eigenbasis of the Hamming graph allows the mean fitness decay to be broken down into contributions of different frequencies. Substituting Δ = *P* Λ*P* ^∗^ in Eqs (5) and (6) and defining 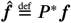 and 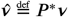 yields:

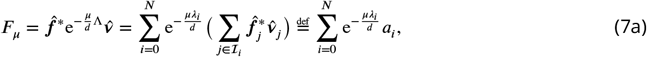

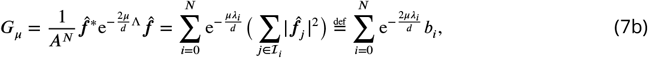

where ℐ_*i*_ groups indices associated with the *i*th eigenvalue and

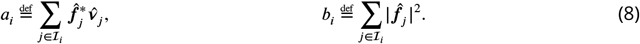

Because *λ*_*i*_ > 0 for *i* ≥ 1, it holds that lim_*μ*→0_ *F*_*μ*_ = *a*_0_ and lim_*μ*→0_ *G*_*μ*_ = *b*_0_. The operation *P* ^∗^***f*** can be understood as a coordinate transformation and the vector 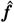 as the landscape ***f*** represented in the Fourier domain. The coefficients *b*_0_, …, *b*_*N*_ are obtained from squaring the modulo of Fourier coefficients and therefore represent the power or energy per frequency *λ*_*i*_. The power spectra of several *NK* landscapes are shown in Fig 6A. For the less rugged landscapes, the power spectrum peaks at a lower frequency than for the more rugged landscapes.

**Figure 6.**
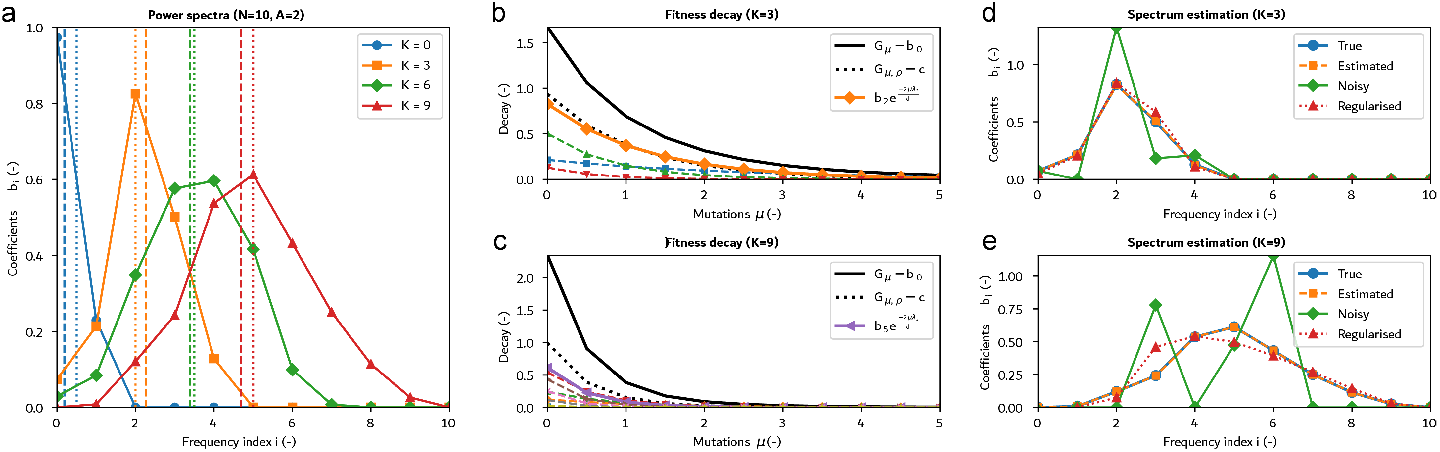
Power spectrum estimation on *NK* landscapes (*N* = 10, *A* = 20). **A:** Power spectra of *NK* landscapes for *K* ∈ {0, 3, 6, 9}. The plot also shows the maximum power frequency from Eq (S2) (dotted) and the frequency obtained from a decay rate based estimation (dashed). **B–C:** Decay rates sampled from the landscapes from Fig 6A for *K* = 3 (top) and *K* = 9 (bottom). The figure shows the decay rate as *G*_*μ*_ − *b*_0_, the fit *G*_*μρ*_ − *c* from Eq (19), and the components *b*_*i*_exp(−2*μλ*_*i*_*/d*) of the five largest *b*_*i*_ of the decay decomposition, with the legend highlighting the dominant one (cf. Fig 6A). **D–E:** Estimation of the full power spectrum via least squares (Eq (15)) and regularised least squares (Eq (17)), with and without noise. The regularisation reduces the impact of noise.

Equations (7a) and (7b) show that as the accumulated mutations *μ* increase, the contribution of each Fourier component to the measured fitness decreases at an exponential rate proportional to *λ*_*i*_, such as shown in Fig 6B. For example, for a linear, zero-mean landscape for which only *b*_1_ ≠ 0, the mean fitness *F*_*μ*_ will decay at a rate proportional to *λ*_1_ = *A* per accumulated mutation, whereas for a maximally rugged landscape for which only *b*_*N*_ ≠ 0, it will decay at a rate proportional to *λ*_*N*_ = *AN*, which is *N* times faster than the linear one. For most landscapes, the decay will be composed from multiple exponentials, such as for the *NK* landscapes in Figs 6B–C.

Although the coefficients *a*_0_, …, *a*_*N*_ and *b*_0_, …, *b*_*N*_ could be estimated from Eq (7a) and Eq (7b), their estimates can become inaccurate (see following section) if *F*_*μ*_ and *G*_*μ*_ are noisy. Assuming that the coefficients *a*_*i*_ and *b*_*i*_ from Eqs (7a) and (7b) vary significantly in magnitude, the fitness decay curve is dominated by the larger-magnitude exponentials. In case of a single dominant component, Eqs (5) and (6) can be approximated by

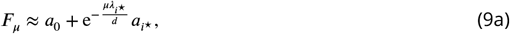

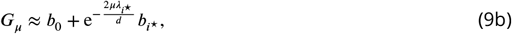

where *i*^⋆^ is the index of the dominant component. Based on Eq (9), we seek to approximate Eq (7a) and Eq (7b) using a single exponential as

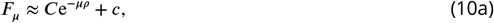

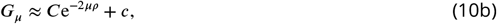

where we used the same variables *C, ρ*, and *c* for *F*_*μ*_ and *G*_*μ*_ although they differ in general. We introduce the decay rate *ρ* as a scalar ruggedness metric that approximates the average of the frequencies *λ*_*i*_ weighted by *a*_*i*_ or *b*_*i*_. This can be shown by expanding Eq (10a) as

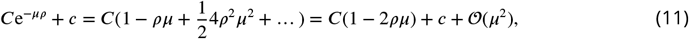

and the right-hand side of Eq (7a) as

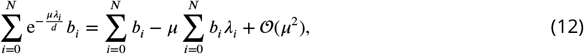

and similarly for Eq (10b) and (7b). Ignoring terms of order 𝒪(*μ*^2^), comparing coefficients yields:

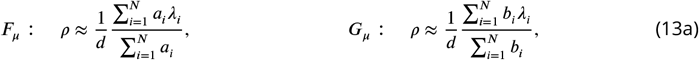

where we used that *c* = *a*_0_ (*c* = *b*_0_) for *F*_*μ*_ (*G*_*μ*_) and 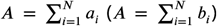, which can be seen from evaluating Eq (10) and (9) for *μ* → ∞ and *μ* = 0. We note that *ρ* for *F*_*μ*_ depends on the initial population ν, whereas for a sufficient large number of samples in Eq (6), *ρ* for *G*_*μ*_ reflects a global property of the landscape. If a single component 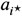 or 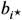 is dominating, then *ρ* is approximately

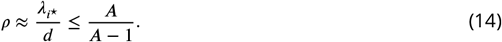

The approximation from Eqs (10a) and (10b) can be analysed in more detail for house-of-cards and *NK* landscapes (see Supplementary, section 4), where it can proved that the approximations hold increasingly well in the limit *N* → ∞ [30].

### Estimating power spectra and ruggedness parameters

Generally, the power spectrum of a fitness landscape is composed from both low-frequency and high-frequency terms, such as shown in Fig 6A. Practically, if the landscape *f* is known, its power spectrum can be computed by reshaping *f* as an *N*-dimensional cube of size *A* along each axis, applying an *N*-dimensional Fast Fourier Transform, and grouping the terms as in Eq (7b).

If the landscape is not known, the power spectral coefficients *b*_*i*_ from Eq (8) can be estimated using the decay curve *G*_*μ*_ from Eq (7b). The procedure for estimating *a*_*i*_ from *F*_*μ*_ is similar. Let 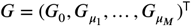 be the squared average fitness measurements of populations accumulating 0, *μ*_1_, …, *μ*_*M*_ mutations and define *b* = (*b*_0_ …, *b*_*N*_)^T^. The following least squares problem can be used to estimate *b* from fitness measurements *G*:

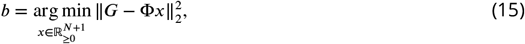

where element *i, j* of the matrix Φ ∈ ℝ^(*M*+1)×(*N*+1)^ is defined as

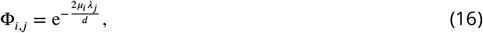

for *i* = 0, …, *M* and *j* = 0, …, *N*. For noise-free observations, problem (15) can be solved using standard techniques to accurately determine the fitness landscape spectrum *b* (Fig 6D–E). However, addition of noise with a signal-to-noise ratio ≤ 10 can already significantly perturb the estimate from Eq (15), which is caused by the ill-conditioned Φ that amplifies the estimation error. To improve the robustness of the estimate, Eq (15) can be extended with a regularisation term as

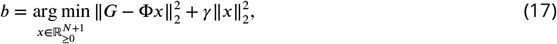

where *γ* ≥ 0 is a regularisation parameter. While the results from Fig 6D–E show that regularisation can improve the estimation accuracy, it depends on having chosen *γ* appropriately; choosing a poor *γ* will lower accuracy. The inference method could be developed further, incorporating more knowledge about the distribution of *b* and the noise process, which is beyond the scope of this paper.

Instead of estimating the full spectrum, we estimate the ruggedness metric *ρ* using the approximations from Eq (10). The scalar parameters *C, ρ*, and *c* are unknown *a priori* and must be estimated from data. As for the power spectrum estimation, the squared average fitness values *G* are recorded for a population accumulating *μ*_1_, …, *μ*_*M*_ mutations. We then seek to find *C, ρ*, and *c* such that

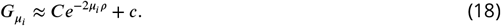

As for the full spectrum estimation, the mutations are stochastic and the decay curves are noisy, which impacts the parameter estimation process. To simplify the estimation, we assume that *G*_0_ is noise free, and that the curve fitted via Eq (18) must pass through it, which allows us to discard parameter *C* by setting *C* = *G*_0_ − *c*. For the remaining data points, we seek to find *ρ* and *c* that minimize the sum of squares error:

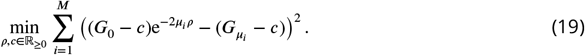

The procedure for estimating *ρ* from *F*_*μ*_ is similar and requires to discard the factor 2 from exponentials. As long as *c* ≠ 0, Eq (19) is a nonlinear optimisation problem that cannot be solved using standard least squares techniques and may have multiple local optima. Here, we are using standard scientific packages for constrained non-linear optimisation (Specifically, the `curve_fit` function from the `scipy.optimize` package [31]), which returns a solution within instants (≤ 1 s) on a standard laptop. Note that the decay rate *ρ* returned by this procedure is generally not of the form in Eq (14), which only admits discrete values as a function of the number of alleles *A* and the protein length *N*. Instead, the *ρ* identified through Eq (19) can take on any intermediate values, capturing a weighted decay rate when several Fourier coefficients are dominant. The curve fitting for two *NK* landscapes is shown in Fig 6B–C, with the estimated decay rates *ρ* shown in Fig 6A. For these examples, the power spectrum peaks at a particular frequency, so that Eq (18) results in a good approximation of the true decay composed from multiple exponentials.

## Code and Data Availability

All code used in this work can be found at GitHub: https://github.com/jessmjames/SLIDE. Data can be accessed from Zenodo: 10.5281/zenodo.16849761.

## Author Contributions

**Sebastian Towers** Responsible for conceptualisation, experimental design, code for simulations, production of figures and writing.

**Jessica James** Responsible for conceptualisation, experimental design, code for simulations, production of figures and writing.

**Harrison Steel** Contributions to conceptualisation and editing.

**Idris Kempf** Project management, contributions to conceptualisation, experimental design, code for simulations, writing and editing.

**Acknowledgment**

IK was supported by EPSRC under the EEBio Programme Grant, EP/Y014073/1.

This preprint was created using the LaPreprint template (https://github.com/roaldarbol/lapreprint) by Mikkel Roald-Arbøl.

