## Supplementary Material for "Sequence-free landscape inference for directed evolution"

### 1 Simulating mutagenesis and selection

In our simulations, we represent a population of $P$ cells at iteration $k$ as $\boldsymbol{p}_k = (\boldsymbol{x}_k^1, \dots, \boldsymbol{x}_k^P)$, where $\boldsymbol{x}_k^i \in \mathcal{G}$ is the genotype of cell $i$. Given an empirical or theoretical landscape $f$, each genotype is associated with a fitness $f_k^i = f(\boldsymbol{x}_k^i)$. When the population $\boldsymbol{p}_k$ is mutagenised, each member $\boldsymbol{x}_k^i$ is exposed to $X$ single mutations to yield $\boldsymbol{x}_{k+1}^i$, where $X$ is an independent identically distributed Poisson random variable. To obtain the decay curves used for ruggedness parameter estimation, we repeat the mutagenesis process for $M$ iterations and record the average population fitness $F_k = 1/P \sum_{i=1}^P f_k^i$, its square $G_k = F_k^2$, and the average accumulated mutations $\mu_k$. This process is repeated and averaged over different starting populations $\mathbf{p}_0$ to generate a decay curve such as the one shown in Fig 3A.

For the selection process used in directed evolution experiments, each iteration is extended with a selection procedure. Here, we use an approach developed in [9] that parameterises the selection using probabilistic "selection functions". The population members $\boldsymbol{x}_k^i$ are ranked according to their fitness $f_k^i$. A fraction of cells is then randomly selected from $\boldsymbol{p}_k$ according to the probability distribution shown in Fig 1 (top right). This distribution is parameterised by two parameters: the base chance $b$ and threshold $t$. If a cell's fitness $f_k^i$ is in the top $t$ percentile (within the population $\mathbf{p}_k$), it is guaranteed to be selected for iteration $k+1$. Otherwise, the selection probability is $b$, which is independent between cells. Here, the fraction of cells selected at each iteration is kept constant as $\mathrm{f} = 0.2$, fixing $t$ as $t = (1-\mathrm{f})/(1-b)$. Each cell that is not selected is replaced by a new cell with genotype taken at random (uniformly and independently) from the pool of cells that were selected. For example, for $t = 0.8$ and $b = 0$ the top $20\,\%$ of fittest cells are selected for the next iteration. The case $t = 1.0$ and $b = 0.2$ represents a selection-free scenario, with a random $20\,\%$ of cells surviving each generation. All simulations were carried out using a GPU-based directed evolution package published in [9].

### 2 Simulating directed evolution

For simulating directed evolution experiments, the selection and mutagenesis procedures are repeatedly applied to a to starting population $\boldsymbol{p}_0$. The selection function is parameterised by the base chance $b$, while the rank threshold $t$ is fixed via $t = (1-\mathrm{f})/(1-b)$. The fraction of selected cells $\mathrm{f}$ is set to $f = 0.2$, so that at each iteration, approxmately $20\,\%$ of the population is selected for the following iteration. The value of $b$ is determined using the decay rate $\rho$. Before starting directed evolution, the population $\boldsymbol{p}_0$ is repeatedly mutagenised without selection for $M$ iterations. The resulting decay curve is used to estimate $\rho$, which is in turn used to select $b$ from a pre-computed map that connects $\rho$ to optimal selection parameters. This map is pre-computed according to the work presented in [9]. Next to the parameter $b$, the work from [9] also incorporates a population splitting parameter $s$ that subdivides the population $\boldsymbol{p}_0$ into $s$ smaller populations, resulting in $s$ smaller directed evolution experiments running in parallel. This is illustrated in Fig 1 (bottom right).

The map linking $\rho$ to the optimal parameter set $(b_\rho, s_\rho)$ is computed using $NK$ landscapes, which relate $\rho$ to $N$ and $K$ via Eq (S2), $\rho = (K+1)/N$. For fixed $N$ and $A$, we generate a number of random $NK$ landscapes for $K = 1, \ldots, N-1$. For each $K$, we run directed evolution experiments for different starting populations $\boldsymbol{p}_0$ and different parameter sets $(b, s)$, and store the best performing parameters for each $K$, defining a map from $(N, K)$ to $(b, s)$. Given $N$ and the estimated $\rho$, the parameters are chosen from the map for which the value $(K+1)/N$ is closest to $\rho$. An example is shown in Fig 1 (middle) for $N = 4$, $A = 20$, and $K \in \{1, 2, 3\}$ ($\rho \in \{0.5, 0.75, 1\}$). Note that while the decay rate $\rho$ can take on continuous positive values, the ruggedness of $NK$ landscapes defined as $\rho_{NK} = (K+1)/N$ can only take on discrete values for fixed $N$. Alternatively, the generated maps can be averaged across different values of $N$.

### 3 Other Ruggedness Metrics

Various measures have been proposed to quantify landscape ruggedness, most of which require both genotypic and phenotypic information. For our analysis, we compare the decay rate measure $\rho$ with five other measures, summarised below. Most metrics were originally defined on an allele set of size 2 ($|\mathcal{A}| = 2$), and are extended here to larger sets ($|\mathcal{A}| > 2$). For a more comprehensive discussion of ruggedness measures, we refer the reader to [7]. We also provide the code used to calculate these metrics at https://github.com/jessmjames/SLIDE.

#### Roughness to slope ratio ($r/s$):

Given a genotype $\boldsymbol{x} = (\sigma_1, \ldots, \sigma_N)^\mathsf{T} \in \mathcal{G}$, a landscape $f_{lin} : \mathcal{G} \to \mathbb{R}$ is *linear* if the contribution of each locus and allele is additive, i.e. $f_{lin}(x)$ can be written as $f_{lin}(\boldsymbol{x}) = \sum_{i=1}^{N} \phi_i(\sigma_i)$, where $\phi_i : \mathcal{A} \to \mathbb{R}$ is the fitness contribution of locus $i$ for allele $\sigma_i$. The roughness to slope ratio [3] captures how well a given landscape $f$ can be approximated by a linear one: After finding the best linear fit $f_{lin}$ to $f$, i.e. the $N \times A$ values $\phi_i(\sigma)$, the mean slope is defined as $s \overset{\text{def}}{=} \sum_{i=1}^{N} \sum_{\sigma \in \mathcal{A}} |\phi_i(\sigma_i)|/(AN)$, the roughness as $r \overset{\text{def}}{=} (\sum_{x \in \mathcal{G}} (f(\boldsymbol{x}) - f_{lin}(\boldsymbol{x}))^2 / A^N)^{1/2}$, and the roughness to slope ratio as $r/s$. For a linear landscape, $r/s = 0$, whereas for more rugged landscapes $r/s$ increases.

#### Landscape $R^2$ measure ($R^2$):

The landscape $R^2$ measure was originally defined in [7] using landscape Fourier analysis and is directly related to the measure $\rho$ introduced in this work. Suppose that the landscape is represented in vector notation as $\boldsymbol{f} \in \mathbb{R}^{A^N}$ and mapped to the Fourier domain as $\hat{\boldsymbol{f}} = \boldsymbol{P}^* \boldsymbol{f}$. Define $b_i \stackrel{\text{def}}{=} \sum_{j \in \mathcal{I}_i} |\hat{\boldsymbol{f}}_j|^2$, where $\mathcal{I}_i$ refers to the indices of $\tilde{f}$ associated with the $n$th eigenvalue of the adjacency matrix. The $R^2$ measure is then defined as $R^2 \stackrel{\text{def}}{=} (\sum_{i=2}^N b_i)/(\sum_{i=1}^N b_i)$. For a linear landscape, $R^2 = 0$, whereas for a more rugged landscape $R^2$ increases. The $R^2$ measure can be computed from decay curves if the full spectrum is estimated via Eqs (15) or (17). However, the difficulties in estimating the full spectrum can make the $R^2$ measure less robust than the decay rate $\rho$. We refer to this quantity as "$R^2$" due to it being equivalent to the Coefficient of determination.

#### Local epistasis ($n_\epsilon$):

Epistasis occurs when the fitness contribution of one of the $N$ loci depends on the allele set at other loci, or in terms of mutations, when the effect of one mutation depends on the presence of others. Simple sign epistasis occurs when a mutation is beneficial in the presence of one set of mutations but disadvantageous in another. For example, consider a landscape with $N = 2$ and $\mathcal{A} = \{0, 1\}$, so that $\boldsymbol{x} = (\sigma_1, \sigma_2)^\mathsf{T}$ with $\sigma_i \in \mathcal{A}$. Simple sign epistasis occurs if changing $\sigma_1$ from $0$ to $1$ has a positive effect on fitness when $\sigma_2 = 0$ and a negative effect when $\sigma_2 = 1$. Here, we measure local epistasis as the number of genotype pairs with a Hamming distance $2$ that exhibit simple sign epistasis.

#### Distance to local maxima ($d_{max}$):

A local maximum is defined as a genotype with higher fitness than all its neighbours. The metric $d_{max}$ measures [5] the Hamming distance to the closest local maximum, averaged over all genotypes. A large $d_{max}$ indicates less local maxima, hence a smoother landscape.

#### Number of direct paths to global maximum ($n_{max}$):

Given a global fitness maximum at genotype $\boldsymbol{x}^\star$, this metric [6] counts the number of fitness-increasing direct (length N) paths $n_{max}$ from the antipodal genotype $\bar{\boldsymbol{x}}^\star$, to $\boldsymbol{x}^\star$. The antipodal genotype $\bar{\boldsymbol{x}}^\star$ is the genotype where, for each locust, the corresponding alleles in $\bar{\boldsymbol{x}}^\star$ and $\boldsymbol{x}^\star$ are different. For $|\mathcal{A}| > 2$, there are multiple antipodal genotypes, so we define $n_{max}$ to be mean over all $(A-1)^N$ possible $\bar{\boldsymbol{x}}^\star$. A small $n_{max}$ indicates a more rugged landscape.

### 4 Special cases for decay rate metric

#### House-of-cards landscapes:

For zero-mean house-of-cards landscapes, the fitness values are assumed to be random, independent, and normally distributed as $\boldsymbol{f}_i \sim \mathcal{N}(0, 1)$. Due to the orthonormality of $\boldsymbol{P}$, this results in all $\hat{\boldsymbol{f}}_i$ being independent of one another, and $\hat{\boldsymbol{f}}_i \sim \mathcal{N}(0, 1)$, i.e. the Fourier representation inherits the same distribution. As a result, the terms $b_i$ in Eq (7b) are chi-squared distributed with mean equal to the multiplicity of $\lambda_i$. The eigenvalue multiplicity $\binom{N}{i}(A-1)^i$ is maximised (The function $\binom{N}{i}(A-1)^i/A^N$ can be interpreted as a binomial distribution, in which case $|i - i_c^\star|/N \leq 3\sqrt{A-1}/(\sqrt{N}A)$ with $i_c^\star = N(A-1)/A$ contains $\sim 99.7\,\%$ of its weight). for $i^\star = \text{round}(N(A-1)/A)$, so that $|b_{i^\star}| \gg |b_i|$ for $i \neq i^\star$. Substituting the eigenvalue $\lambda_{i^\star} = Ai^\star$ yields the decay rate for house-of-cards landscapes:

$$\rho_{HoC} = \frac{1}{d} \times \lambda_{i^\star} = \frac{1}{N(A-1)} \times A\frac{N(A-1)}{A} = 1. \tag{S1}$$

Eq (S1) could also be derived differently. Since every fitness value is independent, with an average fitness of $0$, every time a cell mutates its new fitness is on average $0$. Hence the fitness curve measures how many non-mutated cells are in the population, which is equal to $e^{-\mu}$.

#### $NK$ landscapes:

This analysis can be extended to $NK$ landscapes, which can be viewed as a sum of $N$ smaller house-of-cards landscapes of (effective) dimensionality $K+1$ [30]. For each of the smaller landscapes, the eigenvalue multiplicity is maximised for $i^\star = \text{round}((K+1)(A-1)/A)$, which substituted in Eq (14) yields

$$\rho_{NK} = \frac{1}{N(A-1)} \times A\frac{(K+1)(A-1)}{A} = \frac{K+1}{N}. \tag{S2}$$

Since for $NK$ landscapes the parameter $K \leq N-1$ measures the number of epistatic interactions and therefore ruggedness, Eq (S2) shows that the decay rate definition of ruggedness is closely related to the $NK$ notion of ruggedness.

### 5 Empirical landscapes and ruggedness

To interpret the quantitative predictions of each ruggedness metric it is necessary to consider two factors: the intrinsic properties of the protein and the assay used to generate fitness scores. While we speculate here on differences between landscapes and potential causes for their relative ruggedness, we caution that such interpretations require validation through deeper analysis and complementary measurements to disentangle biological from assay-based contributions.

#### TEV:

The TEV protease (the most rugged in all metrics) recognises a highly specific substrate sequence, and its activity depends acutely on maintenance of its catalytic triad [32], as well as substrate-binding pocket and global fold. Catalytic triads in particular are among the least mutation-tolerant motifs in biology - even single substitutions can abolish function - which may contribute to the highly rugged landscape observed. The assay used to collect this data was based on DNA recording with a base editor [23]. This data is inherently noisy due to the added layer of biological complexity, which is likely to contribute significantly to the ruggedness of this landscape. However, of note is that this assay is much more linearly proportional to activity than the growth-based assays used to measure TrpB and ParD3.

#### ParD3:

ParD3 is an antitoxin in which the C-terminal region is intrinsically disordered in the absence of its toxin partner. Such disordered regions typically rely on general physicochemical properties rather than rigid 3D structures, allowing greater mutational tolerance and producing a smoother landscape [33]. From the assay perspective, this fitness landscape was mapped using a growth competition assay [21]. Such an assay is inherently less noisy than that used to evaluate TEV, due to the fact that the measurement is population level, and the exponential nature of growth also acts to amplify signal at the upper end of fitness. Similarly, growth-based assays of this kind may exhibit "saturation" in the sense that ParD3 variants with toxin binding ability above some threshold at which they can sequester the vast majority of toxin in a cell can thus yield similar (growth-based) fitness scores, even if *in practice* their binding affinity (if it were directly measured) differs significantly. This factor, together with the noise-smoothing properties of this assay, may also contribute to the smoothness of the ParD3 landscape.

#### TrpB:

Similarly to ParD3, TrpB was evaluated using a growth based-assay [22], allowing us to more confidently compare the two on a biological basis. TrpB is a subunit of tryptophan synthase, which catalyses the synthesis of the amino acid tryptophan. Unlike ParD3, it does not contain any large, intrinsically-disordered regions, and thus many residues may be coupled to maintain the folded structure and activity, contributing to epistasis and therefore landscape ruggedness. Meanwhile, in the case of TrpB this growth coupling is achieved through Tryptophan synthesis which, following a similar argument to above, may "saturate" in its impact on measured fitness for high performing TrpB variants (i.e. once an adequate quantity of Tryptophan is produced in the cell further increase in enzyme efficiency is not rewarded by this assay).

### GB1:

GB1 is an immunoglobulin binding protein, whose structure, similarly to TrpB, is globular. Therefore it must also maintain the amino acid interactions that help support folded structure and activity, and consequently contribute to landscape ruggedness. The fitness values in this case were measured using a high-throughput binding assay [2]. The assay achieved a coverage of approximately 100 reads per variant, helping to reduce the noise of the data and therefore assay-related ruggedness, particularly compared to the TEV example.

Given the above arguments, we believe that the ordering of these landscapes by $\rho$ (Fig 4c) is justifiable. The assay used to measure TEV fitness was inherently noisy, alongside the fact that the presence of a catalytic triad in the enzyme could add considerable interaction effects and therefore ruggedness. For this reason, it makes sense that it would be positioned above and not with (as it was by local epistasis, paths to max and distance to local max) the rest of the landscapes - and indeed *all* ruggedness metrics investigated agreed with this conclusion. GB1 and TrpB, both globular proteins that were measured with less noisy assays, have justification to be grouped together (as they were by $\rho$, as well as roughness to slope and landscape $R^2$) and similarly to be below TEV. ParD3 was positioned at the bottom by all metrics, which aligns with our expectations given both its intrinsically disordered structure and the properties of the growth-based assay used to define its fitness.

Considering again Fig 4c and the metric comparisons in Fig 3e we can see significant qualitative differences between metrics. In particular, from Fig 4c the metrics considered can broadly be classified into three groups; Group 1 (decay rate $\rho$, roughness to slope, and landscape $R^2$) place TEV as the most rough, ParD3 as the least, and then group GB1/TrpB close together at intermediate roughness. Group 2 local epistasis and distance to local max) place ParD3 at the bottom, GB1 at intermediate ruggedness, but then group TrpB with TEV at the roughness maximum. Group 3 (paths to max) in turn places all of TEV/GB1/TrpB at the scale's maximum ruggedness - perhaps not surprising because this metric (as shown Fig 3e) saturates very quickly as a landscape becomes more rugged. Considering the biological reasoning above we believe our metric (or more broadly those in Group 1) most aligns with the desirable properties of a ruggedness metric (i.e. as a measure of how sensitive a protein's fitness is to mutation): TrpB with its smoothed growth-based data falls below TEV, as too should GB1. An open question remains whether GB1/TrpB *should* exhibit a similar ruggedness value (as seen for all Group 1 metrics) - this would best be answered with further experimental measurements or augmentation of our comparisons with additional protein landscapes as they become available in the future.
